# Complex trait responses to complex environments: how do larval amphibians navigate co-occurring ecological demands that influence the same traits?

**DOI:** 10.64898/2026.04.24.720614

**Authors:** Stephanie Bristow, Samantha M. Skerlec, Wyatt Mills, Adam Rogers, Amani Saber, Krista J. Ward, Thomas M. Luhring

**Author notes:** **Correspondence:** Stephanie Bristow.

## Abstract

1. Many organisms alter phenotypically plastic traits in response to environmental cues to match their phenotypes with variable environments. In larval amphibians, development and growth rates respond to spatiotemporally variable mortality risk from predation, wetland drying, or resource limitation. However, these rates are also temperature-dependent for ectotherms. Although wild animals experience these factors simultaneously (e.g., thermal regimes, predation risk, resource limitation), most studies investigate their impacts in isolation, limiting our understanding of how they interact across ecological contexts.
2. Here we simultaneously exposed larval Plains Leopard Frogs (*Lithobates blairi*) to varying resource levels and predation risk treatments across a thermal regime to investigate the joint effects of these ecological drivers on growth and development rates and their consequences for size and vagility after metamorphosis. We crossed two predation treatments (waterborne cues from *Procambarus gracilis* fed *L. blairi* larvae, control water) with three food resource levels (5%, 25%, 50% of body mass) and six thermal regimes (diel ± 3°C cycles of 15, 20, 22, 24, 26, 28°C), replicating each combination five times for a total of 180 individuals. We recorded growth and development rates and completion of metamorphosis, then measured juvenile body size and jumping performance.
3. The number of larvae completing metamorphosis was primarily determined by temperature and temperature-dependent effects of resource limitation. Percent metamorphosis peaked at intermediate temperatures when resources were high and were higher in predation-risk treatments at the warmest temperatures. Under high resources, development and growth rates showed unimodal thermal responses that were absent when resources were constrained. Higher resources increased development rates, but proportional increases in growth maintained constant body size across temperatures. Post-metamorphic body size differed only by predation treatment, with predator-exposed individuals being smaller. Juvenile jumping performance increased with body size and individuals raised with high resources without predator cues exhibited the highest performance.
4. The absence of temperature effects on size at metamorphosis reflected unexpected coupling of growth and development rates across treatments, producing uniform body sizes. This pattern contrasts with the temperature-size rule and suggests that plastic responses may exhibit selection for a minimum viable size at metamorphosis.

## INTRODUCTION

Organisms experience the impacts of global climate change within the ecological contexts and constraints of their environment (Masson-Delmotte et al., 2021; Monaghan, 2008; Pascual et al., 2022). Ecological contexts modify the effects of temperature on physiological and life-history processes, altering the magnitude and direction of plastic responses that determine organismal performance (Grigaltchik et al., 2016; Luhring et al., 2018, 2019; Luhring & Delong, 2016; Thomas et al., 2017). Temperature sets the pacing of biological rates (Brown et al., 2004; DeLong et al., 2017; Gibert, 2016; Gillooly et al., 2002) and body size similarly sets constraints on how organisms interact with each other and their environment (Cabrera-Guzmán et al., 2013; Calder, 1984; Peters, 1983; White et al., 2007). Ectotherm body size is shaped by thermal regimes experienced in early ontogeny (Atkinson, 1994) through simultaneous changes in temperature-dependent growth and development rates (Forster & Hirst, 2012). However, body size is not solely determined by temperature. Environmental cues such as resource availability and predation risk exposure in early ontogeny readily alter growth and development rates (Blanckenhorn, 1998; Harkey & Semlitsch, 1988; Lent & Babbitt, 2020; Luhring et al., 2019; Riessen, 1999; Steiner & Van Buskirk, 2008). Despite robust investigations of size-temperature relationships (e.g., Forster et al., 2012) and longstanding appreciation for the impacts of environmental cues in early ontogeny, much less is known of how temperature and environmental cues interactively shape growth rate, development rate, and subsequent body size.

Any factor affecting growth rate or development rate alters an organism’s body size unless these rates increase or decrease simultaneously (i.e., are “coupled”) (**Fig. 1**). It is the decoupling of these rates from each other across a thermal gradient (Forster et al., 2011) that leads to the Temperature Size Rule (TSR) whereby ectotherms mature at smaller body sizes at warmer temperatures (Atkinson, 1994, 1995). Resource availability sets an energy budget for developing organisms, which regulates development and growth rates and constrains phenotypically plastic responses to additional stressors (Courtney Jones et al., 2015; Glazier & Calow, 1992; Steinwascher, 1981). In addition to constraints imposed by resources, warmer temperatures accelerate energetic demands (Schulte, 2015), likely restricting allocation of resources to development and growth at high temperatures.

**Figure 1.**
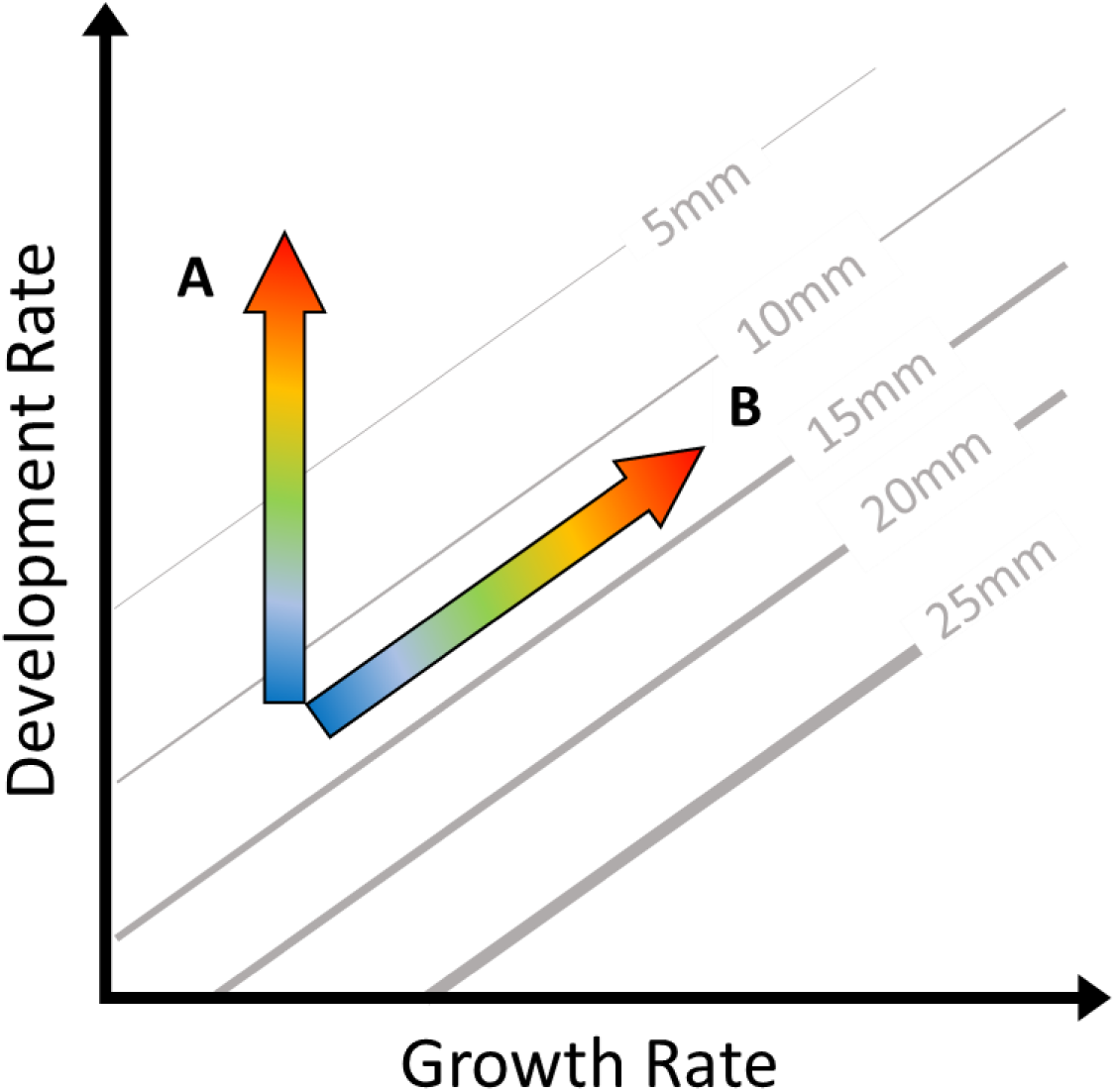
Growth rate (size gained per unit time) and developmental rate (1/time to developmental milestone) determine the body size of an individual at a given stage of development. Grey isoclines indicate combined values of growth and developmental rates that lead to the same size (mm). Temperature effects on growth and development rates influence relative body size (color gradient arrows indicate changes in rates as temperature increases from cold – blue, to warm – red). (A) Increasing development rates and constant growth rates (rates are decoupled) with warming would lead to an increasingly smaller body size at metamorphosis. (B) Size could also remain constant with warming if development and growth rates simultaneously increase (rates are coupled).

Predation risk is experienced in most if not all ecosystems, and many prey species express inducible defenses to increase chances of survival. Antipredator traits include structural defenses and larger bodies to thwart gape-limited predators (Maher et al., 2013; Mori et al., 2005; Riessen & Gilbert, 2019; Urban, 2008; Urban et al., 2017), and accelerated development rates to increase the probability of reproducing (Riessen, 1999), or metamorphosing into a terrestrial landscape to escape aquatic predators (Werner, 1986). Other prey exhibit plastic behavior such as reduced foraging activity around predators to reduce predation risk (Burraco et al., 2013). Predator-induced defenses may increase survivorship but can ultimately alter size at a given developmental milestone (Steiner, 2007; Ward et al., 2023). However, the ability of organisms to respond to predation risk likely depends on existing environmental conditions and energetic demands (Burraco et al., 2013; Costa & Kishida, 2015; Katzenberger et al., 2014; Mauro et al., 2022; Relyea, 2002).

Our focus on identifying causal mechanisms often (and understandably) leads ecologists to assess phenotypically plastic traits individually and in isolation from realistic ecological contexts. There are many excellent mechanistic studies on single traits conducted at “room temperature” with unlimited resources in the absence of predators. However, organisms *in situ* exist under broader thermal regimes and ecological constraints such as predation risk and resource limitation. Warming and predation risk are both known to accelerate development rates, resulting in smaller and younger individuals at developmental milestones (Atkinson, 1995; Beck & Congdon, 2000; Relyea & Hoverman, 2003; Saeed et al., 2021). More recent experiments have demonstrated that responses to predation risk are temperature-dependent themselves (Costa & Kishida, 2015; Luhring et al., 2018, 2019; Padfield et al., 2020; Polo-Cavia et al., 2017). It remains unclear whether metabolic demand increases with the number of experienced stressors (Sokolova & Francisco, 2013), but some studies suggest that resilience to coinciding stressors, such as predation risk or warming, decreases under resource limitation (Bennett et al., 2013; Buskirk, 2000; Steiner, 2007). Temperature, bottom-up, and top-down forces clearly interact to determine ectotherm body size (Bennett et al., 2013; Costa & Kishida, 2015; Huey & Kingsolver, 2019; Zhu et al., 2023). However, joint responses of growth and development rates to multiple ecological variables have not been thoroughly investigated.

In this experiment, we quantified survival, development rate, growth rate, resulting body size and post-metamorphic performance of *Lithobates blairi* (Plains Leopard Frog) across a fully factorial combination of temperature, predation risk, and resource availability treatments. This design allowed us to test how temperature-dependent vital rates jointly respond to bottom-up and top-down ecological pressures. We predicted that development and growth rates would exhibit unimodal thermal responses that would be amplified under high resource availability. We further predicted that predation risk would decouple growth and development rates, leading to smaller body sizes at metamorphosis. Consistent with the temperature-size rule, we expected body size to decline with increasing temperature (**Fig. 1**; Forster et al., 2012). Finally, we predicted that warming, predation risk, and resource limitation would each reduce juvenile body size and locomotor performance.

## METHODS

### Study species

Our study species, *Lithobates blairi* (Plains Leopard Frog), is distributed throughout the Midwestern United States and belongs to a cosmopolitan group of “leopard frogs” common across North and Central America (Gibbons et al., 2006; Giovanelli et al., 2008). Many temperate amphibians breed in ephemeral pools (Wilbur, 1980) which experience high diel temperature variation and seasonal drying (Burkhead et al., 2022; Calhoun & DeMaynadier, 2007; Zedler, 2003). We selected *Procambarus gracilis (*Prairie Crayfish) to produce predation risk cues because they are natural predators of amphibian larvae (Gherardi et al., 2001) and occur in pools used by *L. blairi* at our collection site.

### Species collection

Animal collection and handling procedures were approved by the Wichita State University Institutional Animal Care and Use Committee (IACUC; protocol #283a) and conducted in accordance with institutional and national guidelines. *Lithobates blairi* egg masses were collected from Wichita State University Biological Field Station: Youngmeyer Ranch (YMR). YMR is an active cattle ranch in the Flint Hills region of Kansas characterized by native tallgrass prairie with ephemeral wetlands and intermittent streams (Burkhead et al., 2022; Houseman et al., 2016). To incorporate local genetic variation, seven egg masses were collected from a YMR wetland (37.33400 N, −96.30023 S) on 31 March 2022.

Egg masses were refrigerated overnight before transport to outdoor mesocosms at Wichita State University’s Biological Field Station: Ninnescah Reserve, Viola, KS the next day. Eggs were then divided among five, 1,000 L holding tanks prepared with 110 L soil from an on-site ephemeral pool, and 300 g dried wetland vegetation (see Ward et al. 2023; Skerlec & Luhring 2023).

Eggs hatched on 10 April 2022, and larvae were allowed to reach Gosner stage 25 and a minimum mass of 150 mg, where individuals are free-swimming and easily handled (Gosner, 1960; Schoeppner & Relyea, 2005). On day 0 of the experiment (8 May 2022), approximately 40 larvae from each holding tank were placed into a 5-gallon bucket and allowed to intermix before being randomly selected and assigned to treatment replicates.

Predators used in the study (30 *Procambarus gracilis*) were collected from Chautauqua County, Kansas (37.05797 N, −96.18156 S). Excess *L. blairi* larvae from holding tanks were routinely euthanized and fed to crayfish to produce predation cues (see Experimental design).

### Experimental design

We selected six temperatures (15, 20, 22, 24, 26, 28°C) based on temperature ranges recorded in YMR pools (Burkhead et al., 2022). Six growth chambers (Percival Scientific, model I36VL) were programmed with 24-hour ramping cycles reaching ± 3°C around the mean treatment temperature (e.g., the 15°C chamber reached 18°C at 18:00 and 12°C at 06:00). Lighting followed a 15:9 light:dark cycle to reflect seasonal light availability, while humidity was maintained at 80% to minimize water evaporation.

In aquatic systems, prey commonly respond to chemical predator cues (i.e., ‘kairomones’) and alarm cues released from injured conspecifics (Crawford et al., 2012; Crowl & Covich, 1990; Howe, 1976; Schoeppner & Relyea, 2005, 2009). Predation risk was simulated using chemical cues produced by crayfish feeding on larval conspecifics. Crayfish were individually housed in clear 4.4 L containers with 3.0-3.5 L of natural rainwater. During each 50% water change (three times weekly), crayfish (*N* = 10) were fed ∼300mg of previously euthanized larvae (Relyea, 2006). Within two hours of feeding, water from crayfish containers was filtered through 63μm sieves and combined in a 50 L carboy to homogenize cues (hereafter “predation cues”).

Half of the larvae (*N* = 90) were housed in rainwater with predation cues for the first 20 weeks of the study (and maintained without cues thereafter), while the remaining larvae (*N* = 90) were housed without predation cues. Each replicate consisted of a single larva in an opaque, plastic container (IUMÉ, 769 mL total volume) filled with 500 mL water and covered with a transparent lid containing six air holes (1 cm in diameter). Containers received 50% water changes three times weekly throughout the experiment.

Amphibian larvae experience variable resource conditions *in situ* (Courtney Jones et al., 2015), as primary productivity increases with temperature (Williams et al., 2014) and competition shifts over time (Travis, 1984). Resource treatments were assigned based on body mass: low = 5%, medium = 25%, and high resources = 50%. The high resource treatment approximated *ad libitum* feeding, 5% is known to depress growth rates, and 25% represented an intermediate treatment (Bennett et al., 2013). Larvae were fed three times weekly according to their body mass at the start of each week. Larvae were weighed weekly using a drying device (2.5 cm PVC with window screen), blotted dry before measurement on a digital scale (Ohaus, model NV222).

Food consisted of blended rabbit chow (National Research Council, 1974) mixed with unflavored gelatin and agar to reduce fouling. Initially, larvae received 100 mg of food three times weekly until reaching 200 mg body mass, after which feeding followed assigned resource treatments. Water changes, feeding, and measurements were conducted concurrently to minimize handling time, and each individual was handled <5 minutes per week.

Due to a lack of growth and metamorphosis in low resource treatments, this treatment was removed from analyses. Predation treatments (present or absent) were therefore crossed with two resource levels (medium, & high) across six temperatures (15-28°C). Each combination was replicated five times for a total of 120 replicates included in our analysis.

### Data collection

Data collection took place over 38 weeks (May 2022-February 2023). Percent metamorphosis was calculated as the proportion of replicates completing metamorphosis within each temperature by predation risk by resource level treatment combination. For each individual, we estimated SVL, larval development rate and growth rate up to Gosner stage 38 (pre-metamorphosis), and juvenile SVL and jumping performance after metamorphosis.

Larval SVL and Gosner stages were recorded twice weekly for 12 weeks and weekly thereafter. Individuals were photographed in petri dishes on graph paper using a digital camera (Olympus Tough TG-6 4K, Model IM015). SVL measurements were obtained using ImageJ (Abramoff et al., 2004; Schneider et al., 2012) by drawing a line from the snout to the base of the tail.

### Development & growth rates

Development and growth are relatively linear until Gosner stage 38, before major morphological changes associated with metamorphosis (Grosjean, 2005; Rita et al., 2010). We therefore used Gosner stage 38 to determine larval development and growth rates. Development rate was calculated as 1/days to stage 38, and growth rate as the change in body size over the larval period (ΔSVL/days to stage 38). Pre-metamorphic corresponds to SVL (mm) at stage 38.

### Juvenile size & jumping performance

After reaching Gosner stage 44, water was reduced by 80% to allow individuals to complete metamorphosis. Most frogs completed metamorphosis and resorbed their tails within two days (Gosner stage 46). Jumping performance trials were conducted indoors at 22°C in a walled arena (3 × 0.5 × 0.5-meter runway) lined with damp paper towels on a tarp (Sinsch et al. 2020). Each juvenile was placed in the center of the arena and covered with an opaque container for one minute to acclimate. After removing the container, juveniles were allowed to jump five times, and if an individual paused for >20 seconds, they were prodded gently on the urostyle. Distances between landing positions were recorded, and the average of the three longest jumps was used as an estimate of optimal jumping performance. Juvenile morphometrics were recorded after trials to minimize handling effects on performance.

### Statistical analyses

Thermal performance traits are typically nonlinear across temperatures (Angilletta, 2009; Rebolledo et al., 2021; Rezende & Bozinovic, 2019). Generalized additive models (GAMs; Wood, 2006) incorporate smoothers to account for nonlinear responses across continuous predictors, such as temperature. Therefore, we used GAMs to analyze percent metamorphosis, development rate, and growth rate. Model diagnostics were assessed using gam.check and treatment and smoother effects were evaluated with anova.gam function in the mgcv package (Wood, 2006). Using the glm2 package, generalized linear models (GLMs) were fit to response variables lacking significant temperature smoother terms during preliminary GAM construction. Initial GLM structure included predation risk, resource level, and their interaction, and non-significant interaction terms were removed to obtain the most parsimonious model. ANOVAs were performed via the car package with Type III sums of squares for models with interactions and Type II sums for models without interactions. Analyses were conducted in RStudio version 4.1.1 (R Core Team 2021),

## RESULTS

### Metamorphosis

Thermal extremes and resource limitation successfully captured the full range of tolerated eco-physiological limits. The low resource treatment was removed from this study due to a lack of growth, and only two larvae from 15°C treatments reached metamorphosis by week 38. Data from the 15°C treatments were included for metamorphosis results but was dropped from all other analyses. Nearly three quarters (73%) of larvae (excluding 15°C and low resource treatments) successfully reached metamorphosis. Percent survival to metamorphosis was unimodal with respect to temperature (F = 10.0, P = 0.001) and the shape of that unimodal relationship was further shaped by resource availability (F = 13.0, P = 0.011) (**Table 1**, **Fig. 2**). All curves for percent metamorphosis showed peaks at intermediate temperatures and were higher at warmer temperatures for larvae raised with medium resources. Predation risk also showed a tendency to elevate percent metamorphosis at warmer temperatures (F = 5.9, P = 0.074).

**Figure 2.**
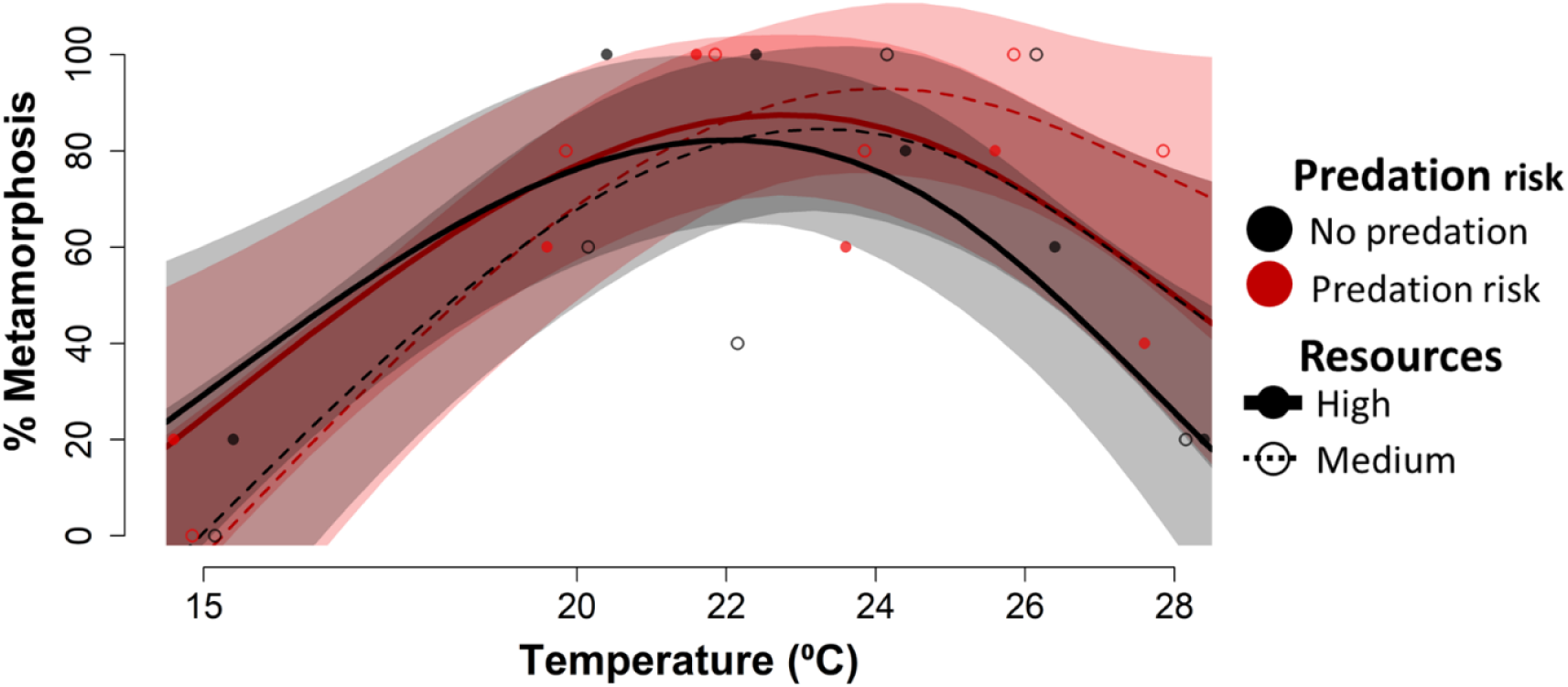
Relationship between larval *Lithobates blairi* percent metamorphosis and the combined effects of resource availability, predation risk and temperature. Points represent raw observations, with color indicating predation treatment (black = no predation risk; red = predation risk) and symbol type indicating resource level (filled circles = high resources, open circles = medium resources). Lines depict generalized additive model (GAM) fits for each treatment combination, with solid lines representing high resources, and dashed lines representing medium resources. Shaded regions represent 95% confidence intervals.

**Table 1.** Summary table of parametric (left) and smoother (right) terms from the generalized additive model (GAM) used to analyze proportion of individuals reaching metamorphosis in each predation risk (no predation & predation risk) by resource (high & medium) treatment combination across a 15-28°C thermal regime.

| Response | Parametric Terms | df | F | P | Smoother Terms | Ref.df | F | P |
| --- | --- | --- | --- | --- | --- | --- | --- | --- |
| % Metamorphosis | Predation risk | 1 | 1.002 | 0.331 | **Temperature** | **2.134** | **9.992** | **0.001** |
|  | Resource | 1 | 0.040 | 0.844 | Temperature: No predation | 1.110 | 0.251 | 0.766 |
|  |  |  |  |  | Temperature: Predation risk | 0.625 | 5.859 | 0.074 |
|  |  |  |  |  | Temperature: High resource | 0.625 | 2.056 | 0.274 |
|  |  |  |  |  | **Temperature: Medium resource** | **0.625** | **13.047** | **0.011** |

### Development & growth rates

GAMs were fit to development and growth rates for 20-28°C larvae that successfully reached Gosner stage 38 (*N* = 79). Both physiological rates showed strong temperature-dependent responses under high resource levels (development rate: F = 11.8, P <0.001; growth rate: F = 5.4, P = 0.002) and were relatively temperature invariant when resources were reduced (**Tables 2** & **3**; **Fig. 3**). Predation risk caused temperature-independent (i.e., additive, not interactive) increases in development rate (F = 4.1, P = 0.047) and similar but marginal increases in growth rate (F = 3.2, P = 0.076) (**Tables 2** & **3**). Growth and developmental rates under high resources peaked around 24-26°C.

**Table 2.** Summary table of parametric (left) and smoother (right) terms from the generalized additive model (GAM) used to analyze responses of pre-metamorphic development rates (1/days to Gosner stage 38) across predation risk (no predation & predation risk), resource (high & medium), and temperature (20-28°C) treatments.

| Response | Parametric Terms | df | F | P | Smoother Terms | Ref.df | F | P |
| --- | --- | --- | --- | --- | --- | --- | --- | --- |
| Development rate | Predation risk | 1 | 4.091 | 0.047 | Temperature | 0.500 | 12.548 | 0.015 |
|  | Resource | 1 | 19.802 | <0.001 | Temperature: No predation | 0.933 | 0.818 | 0.385 |
|  |  |  |  |  | Temperature: Predation risk | 0.625 | 0.361 | 0.636 |
|  |  |  |  |  | Temperature: High resource | 2.586 | 11.792 | <0.001 |
|  |  |  |  |  | Temperature: Medium resource | 1.512 | 0.578 | 0.648 |

**Table 3.** Summary table of parametric (left) and smoother (right) terms from the generalized additive model (GAM) used to analyze responses of pre-metamorphic growth rate (mm/day) across predation risk (no predation & predation risk), resource (high & medium), and temperature (20-28°C) treatments.

| Response | Parametric Terms | df | F | P | Smoother Terms | Ref.df | F | P |
| --- | --- | --- | --- | --- | --- | --- | --- | --- |
| Growth rate | Predation risk | 1 | 3.245 | 0.076 | Temperature | 0.500 | 3.479 | 0.191 |
|  | Resource | 1 | 12.140 | <0.001 | Temperature: No predation | 0.625 | 1.863 | 0.284 |
|  |  |  |  |  | Temperature: Predation risk | 0.814 | 0.200 | 0.688 |
|  |  |  |  |  | Temperature: High resource | 2.596 | 5.392 | 0.002 |
|  |  |  |  |  | Temperature: Medium resource | 0.625 | 0.030 | 0.892 |

**Figure 3.**
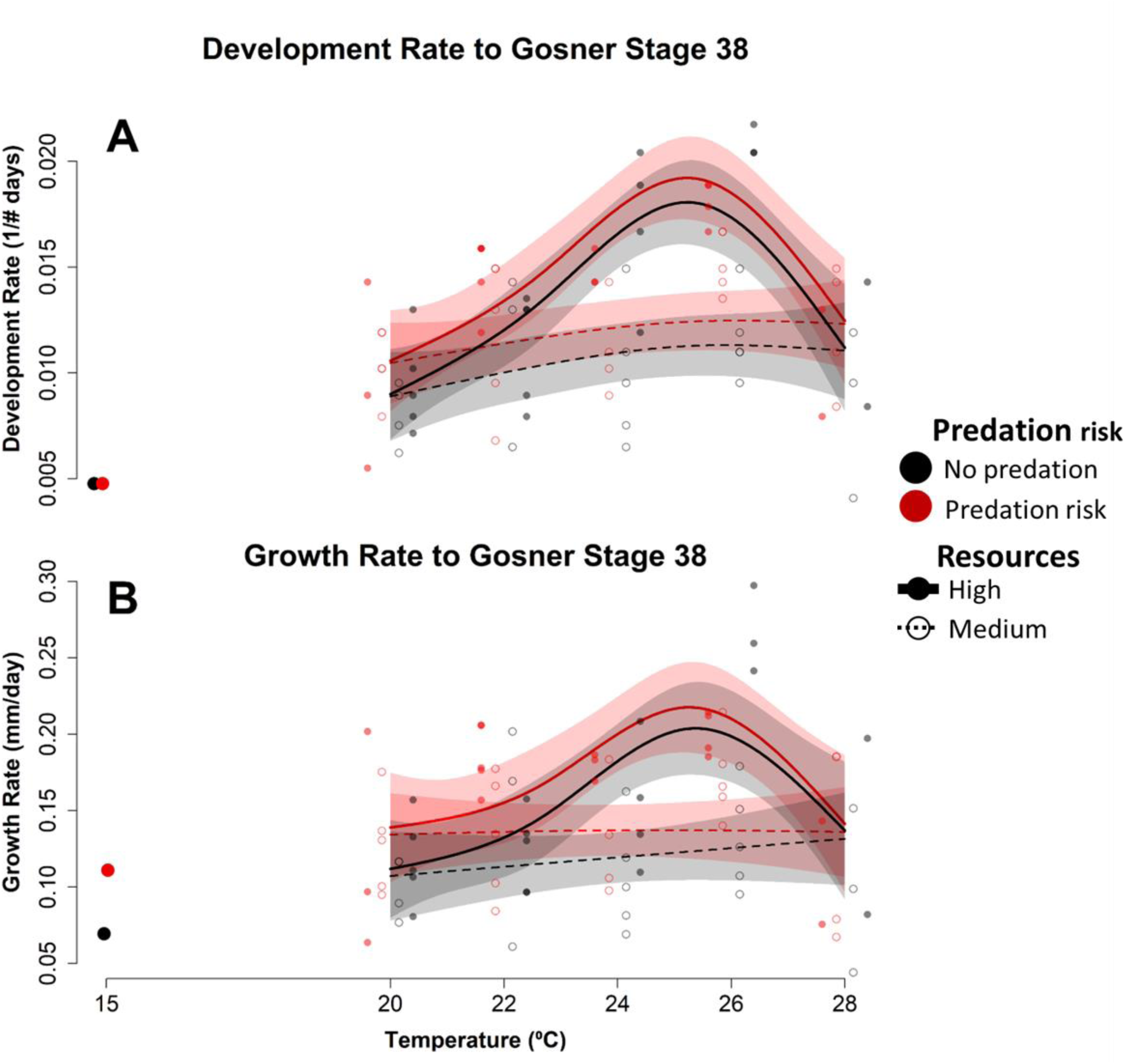
Relationship between larval *Lithobates blairi* **(A)** development rates (1/days to Gosner stage 38) and **(B)** growth rates (mm SVL/day) across temperature, resource availability, and predation risk and temperature. Points represent raw observations, with color indicating predation treatment (black = no predation risk; red = predation risk) and symbol type indicating resource level (filled circles = high resources, open circles = medium resources). Lines depict generalized additive model (GAM) fits for each treatment combination, with solid lines representing high resources, and dashed lines representing medium resources. Shaded regions represent 95% confidence intervals. Development and growth rates for the two individuals at 15 °C that reached Gosner stage 38 are shown as individual points alongside the fitted GAMs (see Results for details).

### Ecological context and temperature shape development, growth, & body size

Development and growth rates remained tightly coupled (i.e., they followed size isoclines **Fig. 1 & 4**) but demonstrated non-linear plastic responses across temperature with interactive effects of resource availability and predation risk. Peaks in growth and development rates at intermediate temperatures (e.g., 24-26°C) led to similarly sized individuals metamorphosing earlier in high resource treatments (moving up the size isocline in **Fig. 4**). Although similar patterns of rate coupling were found in medium resource treatments, ranges of growth and development rates were constrained with substantial overlap across temperatures.

**Figure 4.**
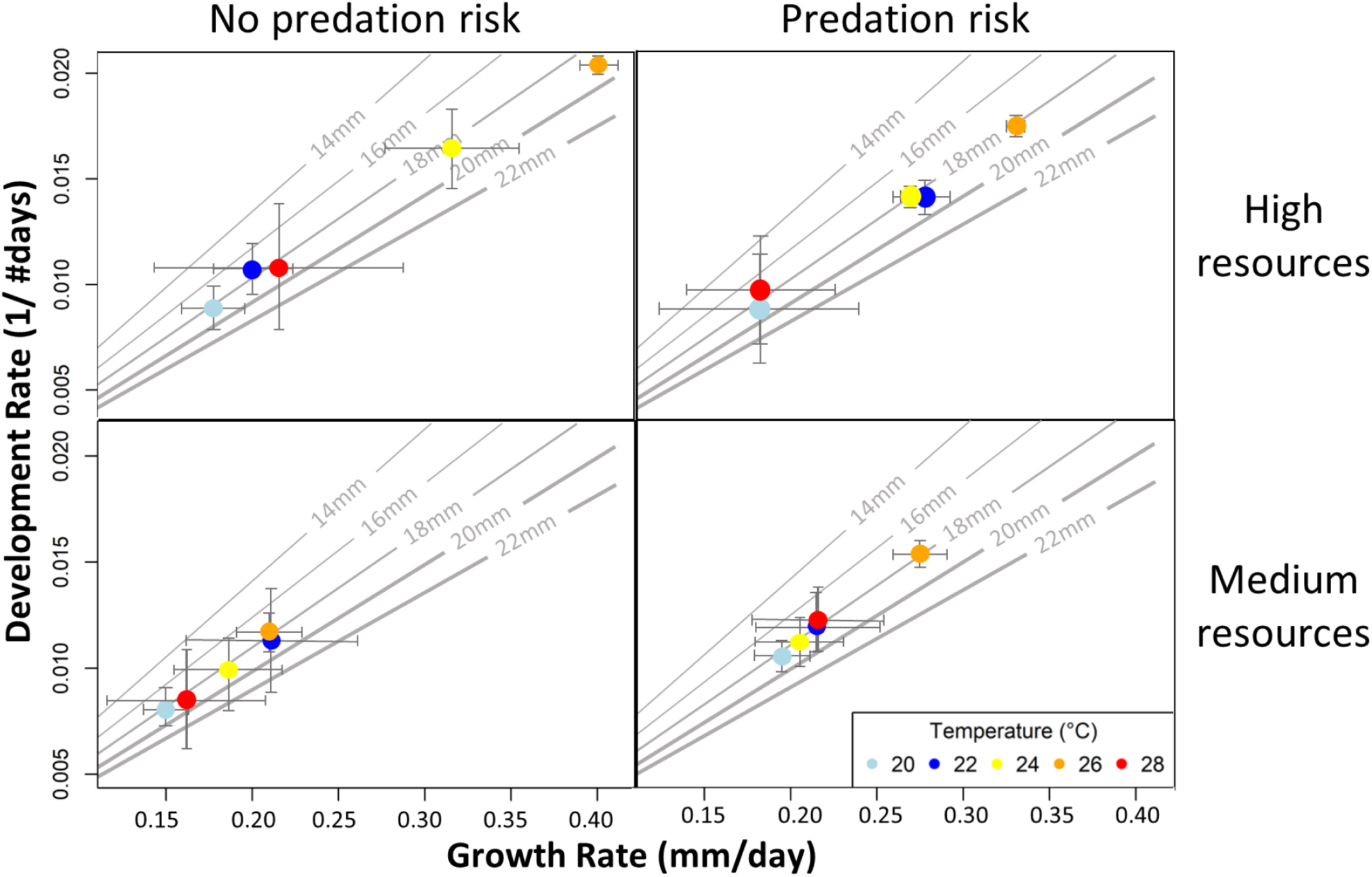
Coupled changes in development and growth rates across treatments. Mean development rate (1/days to Gosner stage 38) is plotted against mean growth rate (mm /day), with both axes jointly determining body size at stage 38 (see Fig. 1 for conceptual framework). Diagonal isoclines represent combinations of development and growth rates that yield equivalent snout vent length (SVL; mm), with thicker isoclines indicating increasing body size. Movement along isoclines reflects trade-offs between growth and development that result in similar final body size. Within each panel, points represent treatment-specific means at each temperature, and 2-dimensional error bars denote 95% confidence intervals for growth and development rates.

### Size pre- & post- metamorphosis

Due to a lack of non-linearity across temperature, SVL measurements were analyzed via GLMs. Larval, or pre-metamorphic SVL (Gosner stage 38) did not significantly change with any treatment (temperature: F = 1.8, P = 0.190 predation risk: F = 0.7, P = 0.412; or resource availability: F = 0.5, P = 0.501). However, juvenile, or post-metamorphic SVL (Gosner stage 46) significantly varied with predation risk treatments. Juveniles from non-predation treatments were 4.05% (95% CI: 0.6-7.5%) larger than juveniles from predation risk treatments (χ^2^ = 5.6, P = 0.018) (**Table 4**, **Fig. 5**).

**Figure 5.**
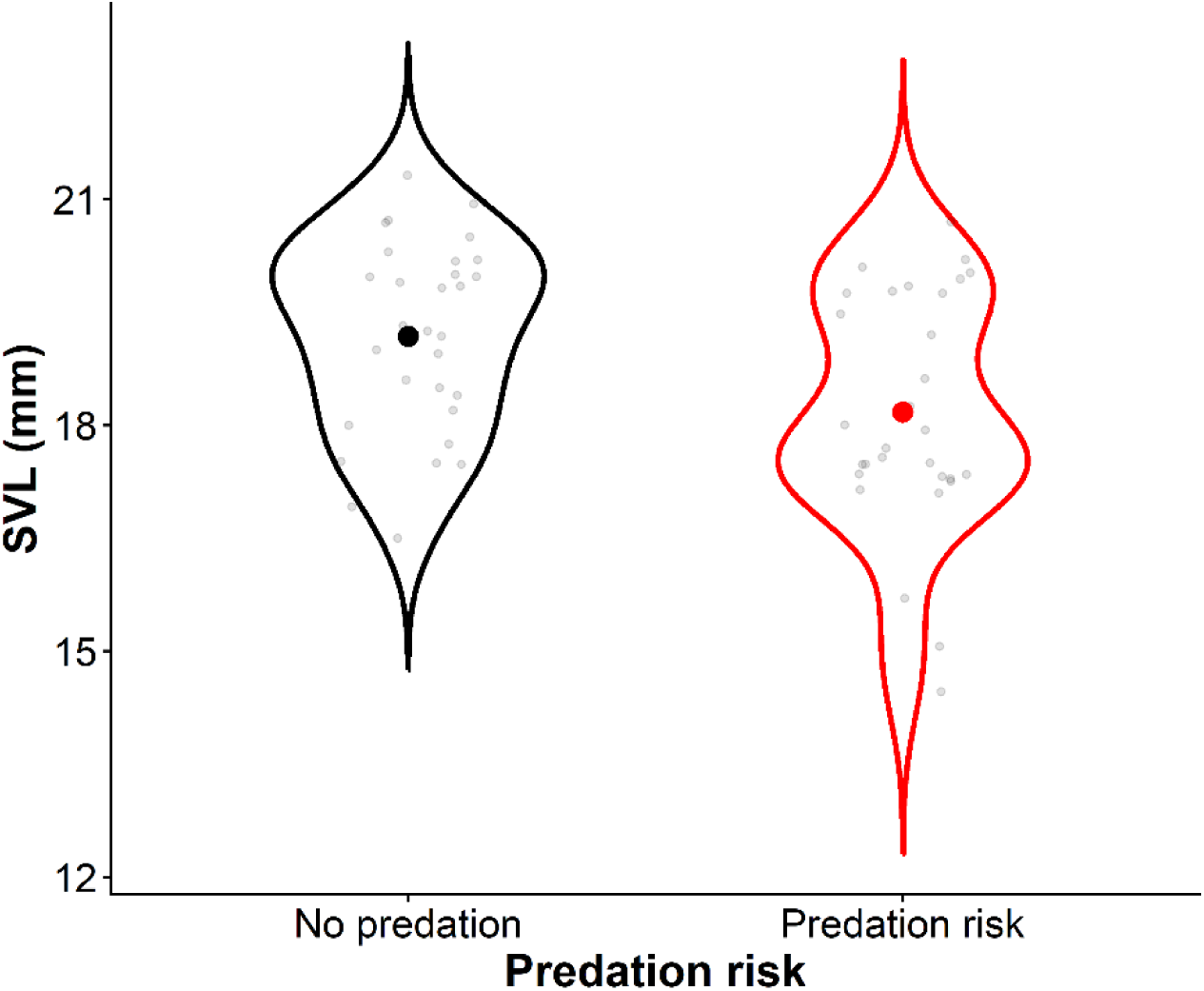
Relationship between juvenile *Lithobates blairi* snout vent length (SVL; mm) and the effects of predation risk. Violins are plotted against raw data points, and circles depict generalized linear model (GLM) mean SVL estimates for each predation risk treatment.

**Table 4.** Summary table of term statistics from the generalized linear models used to analyze responses of pre-metamorphic and juvenile SVL and jumping performance (average distance of 3 longest jumps).

| Response | Parametric Terms | df | $\chi^2$ | P |
| --- | --- | --- | --- | --- |
| Pre-metamorphic SVL | Temperature | 1 | 1.512 | 0.219 |
|  | Predation risk | 1 | 0.613 | 0.434 |
|  | Resource | 1 | 0.458 | 0.499 |
| Post-metamorphic SVL | Temperature | 1 | 1.392 | 0.238 |
|  | **Predation risk** | **1** | **5.618** | **0.018** |
|  | Resource | 1 | 2.602 | 0.107 |
| Jumping performance | SVL | 1 | 24.815 | **<0.001** |
|  | Temperature | 1 | 0.008 | 0.928 |
|  | **Predation risk** | **1** | **5.454** | **0.020** |
|  | **Resource** | **1** | **10.245** | **0.001** |
|  | **Predation: Resource** | **1** | **5.224** | **0.022** |

### Juvenile jumping performance

Larval conditions related to resource availability and predation risk had interactive effects that carried over to jumping performance (χ^2^ = 4.6, P = 0.032) (**Table 4**). This interaction was primarily driven by juvenile size. Juveniles from no predation-high resource treatments jumped farther than any other treatment combination (**Fig. 6**). Even within the high resource treatment, frogs unexposed to predation risk jumped on average 19.5% ± 9.7-29.2% (mean ± 95%CI) farther in their longest three jumps than those exposed to predation risk.

**Figure 6.**
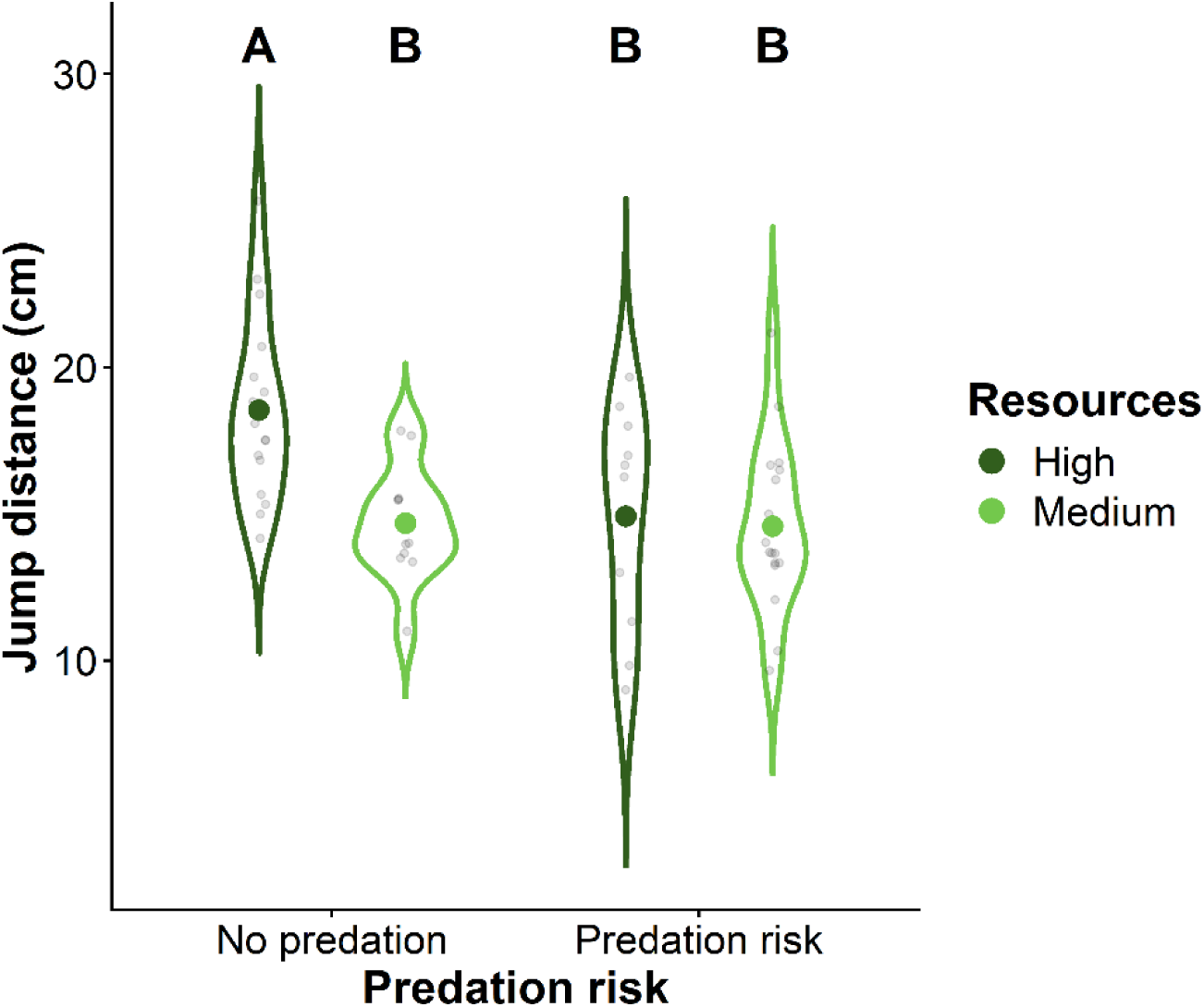
Relationship between juvenile *Lithobates blairi* average jump distance (cm) and the joint effects of predation risk and resource availability. Violins are plotted against raw data points, and circles depict generalized linear model (GLM) mean estimates of jump distance for each treatment combination. Letters depict significant differences between groups based on Tukey HSD pairwise comparisons.

## DISCUSSION

We showed that survival, growth, and development rates were temperature-dependent when resources were abundant but were unexpectedly temperature-invariant when resources were limited. Furthermore, predation risk affected vital rates consistently across temperature (additive effect). Although vital rates changed uniquely across temperature according to different treatments, growth and development rates remained tightly coupled and did not demonstrate the decoupled response predicted by the temperature-size rule (TSR). Predation risk reduced post-metamorphic body size and, together with resource limitation, reduced juvenile jumping performance which was also strongly correlated with body size. Collectively, these findings demonstrate how ecological context, particularly resource availability, governs whether temperature-dependent phenotypic plasticity is expressed and can alter expected relationships between temperature and body size.

### Ecological context modifies temperature-dependent survival & rates

Climate (Lowe et al., 2021) and biotic factors such as predation risk and resource availability (Courtney Jones et al., 2015; Relyea, 2007) strongly influence development and survival. However, most studies examine only one or two ecological drivers at a time and rarely include a full thermal gradient to evaluate temperature-dependent responses (Álvarez & Nicieza, 2002; Bennett et al., 2013; Dastansara et al., 2017; Ohmer et al., 2023). In our study, the proportion of individuals completing metamorphosis exhibited a unimodal response across temperature. However, few individuals raised under low resource conditions reached metamorphosis or even later Gosner stages within the experimental timeframe, reflecting strong energetic constraints on development. This pattern is broadly consistent with previous work showing that survival generally increases with resource levels or reduced competition for resources (Newman, 1998). Interestingly, we found that more individuals completed metamorphosis in medium resource treatments compared to high resource treatments. Reduced survivorship at warmer temperatures under high resource treatments could be driven by increased bacterial growth associated with excess food or waste, which can degrade water quality (Earhart et al., 2022; Frede & Lignell, 1997; Mishra et al., 2022).

Exposure to predation risk marginally increased the proportion of individuals reaching metamorphosis at warmer temperatures. A potential explanation for this pattern is that predation risk may induce the upregulation of heat-shock proteins (HSP) in aquatic organisms (Pauwels et al., 2005; Slos & Stoks, 2008). HSPs are known to support resistance to thermal stress and survival (Gao et al., 2014; Sørensen et al., 2009). Consequently, predator-induced HSP expression has been hypothesized to increase survival under thermal stress (Luhring et al., 2019; Luhring & Delong, 2016; Slos & Stoks, 2008), which could explain the slightly higher survival rates observed at warmer temperatures in our study. However, the direct role of predator-induced HSP expression in improving survival under warming remains largely unknown.

### Growth-development coupling, energetic constraints, & the TSR

The temperature size rule suggests that organisms develop faster but reach a smaller body size at higher temperatures, often due to the decoupling of growth and development rates (Atkinson, 1994; Forster et al., 2012; Perrin, 1995). In contrast, development and growth rates of *L. blairi* in our study were both temperature sensitive (peaking at intermediate temperatures) and tightly coupled across a 13°C thermal gradient. This tight coupling of growth and development rates led to temperature-independent body size at metamorphosis, contradicting the TSR seen in a variety of aquatic ectotherms. Larvae provided with high resources reached faster development rates at intermediate temperatures but were offset by proportional increases in growth rates (Fig. 4). This growth-development coupling allowed individuals to reach metamorphosis earlier without sacrificing body size, suggesting that sufficient resources maintain coupling between growth and development rather than the decoupling predicted by the TSR. This pattern is consistent with studies showing that temperature-dependent effects on growth and body size depend strongly on resource availability in striped marsh frogs (Courtney Jones et al., 2015), and resource quality in the Chinese giant salamander (Zhu et al., 2023).

*L. blairi* in our experiment were smaller than full siblings raised in outdoor mesocosms (Skerlec & Luhring, 2023) which may be related to constraints of a standardized laboratory diet. The resulting narrow size range in our study may have limited our ability to detect treatment effects. Future work is necessary to evaluate how diet quantity and quality interact with temperature and predation risk to affect growth.

### Predation risk coordinating plasticity

Predation risk induces diverse phenotypically plastic responses in early development that increase probability of survival. In many aquatic systems, predators influence prey behavior, with a common response being reduced foraging in the presence of predators and a resulting tradeoff in growth (Arnett & Kinnison, 2017; Bourdeau & Johansson, 2012; Semlitsch & Reyer, 1992; Skelly & Werner, 1990). In our study, predation risk moderately increased development rates in a manner that was independent of temperature. Consequently, predation risk had little effect on body size within the larval stage, even though individuals exposed to predation risk ultimately reached smaller post-metamorphic body sizes. It is possible that predation risk altered energy allocation during development (e.g., retention of lipid reserves) without significantly decoupling growth and development rates. While predator-induced shifts in resource allocation have been documented in some systems (McPeek et al., 2001; Slos & Stoks, 2008; Trussell et al., 2006), further work is needed to determine how prey partition energy among growth, maintenance, and defensive traits when exposed to predation risk. Predation risk and other variables affecting energy allocation may become especially important under climate warming, as higher temperatures elevate metabolic demand, and may amplify energetic tradeoffs during development.

### Carryover effects & fitness consequences

Environmental conditions in early life stages can influence performance in following life stages, yet most studies focus on larval or juvenile traits independently. In our study, jumping performance reflected both juvenile size (which varied with exposure to predation risk) and resource availability provided in the larval stage. Our results suggest predation risk accelerates larval development rates, likely increasing chances of reaching metamorphosis, but these responses may introduce tradeoffs for body size and locomotor performance. These tradeoffs in size could influence fitness in terrestrial environments, as smaller individuals face higher risks with desiccation (Child et al., 2008) and reduced foraging success (Cabrera-Guzmán et al., 2013). Although our findings highlight how larval conditions shape juvenile performance, future work is needed to explore how developmental plasticity affects survival and reproductive success. In particular, compensatory growth following metamorphosis may offset early developmental constraints (Boone, 2005; Székely et al., 2020; Thompson & Popescu, 2021).

### Climate change implications

Climate warming will simultaneously increase metabolic demand and alter resource availability in freshwater ecosystems. Our findings show that resource availability strongly mediates plastic responses to temperature, suggesting that populations facing resource limitation could be more vulnerable to warming effects on development and survival. In our study, larvae maintained similar body sizes rather than following the temperature-size rule, indicating that maintaining a minimum viable size at metamorphosis may override temperature effects on body size. However, if warming increases energetic demand while resources become limited, developmental success and rates of metamorphosis may decline. In addition, warming may increase predator metabolic demand and potentially strengthen top-down pressures on prey. Future research should therefore examine how resource limitation and predation risk during early development interact with warming and whether these developmental conditions generate lasting effects on performance and fitness across life stages.

## Conflict of Interest

The authors declare no conflict of interest in relation to the content or publication of this manuscript.

## Author Contributions

SB and TML conceived and designed the study. SB, SMS, WM, AR, AS, and KJW collected and curated the data. SB analyzed the data. SB and TML wrote the manuscript. All authors contributed critically to the drafts and gave final approval for publication.

## Data Availability

Data and code will be made publicly accessible on Zenodo upon acceptance for publication at https://doi.org/10.5281/zenodo.19207231

## Acknowledgements

Stephanie Bristow was supported by MWPARC, the Jayne & Glenn Milburn Fellowship, and the Claude L. Sheats Jr. Environmental Science Endowed Fellowship. Thomas M. Luhring’s contributions were supported by Kansas NSF EPSCoR. We thank Dexter Mardis for assistance with field collections. We also thank Cindy Gilbert, Clarke Burgert, Corbin Mai, Erin Schaefer, George Kluczykowski, John Suenram, Kayla Brizendine, Lindsey Choi, Makena Frazer, Parker Binns, and William Cook for assistance with data collection.

## Notes

https://doi.org/10.5281/zenodo.19207231

### Competing Interest Statement

The authors have declared no competing interest.

## REFERENCES

Abramoff, M. D., Magalhaes, P. J., & Ram, S. J. (2004). Image Processing with ImageJ. Biophotonics International, 11(7), 36–42.

Álvarez, D., & Nicieza, A. G. (2002). Effects of induced variation in anuran larval development on postmetamorphic energy reserves and locomotion. Oecologia, 131(2), 186–195. 10.1007/S00442-002-0876-X.

Angilletta, M. J. (2009). Thermal Adaptation: A Theoretical and Empirical Synthesis. Oxford University Press.

Arnett, H. A., & Kinnison, M. T. (2017). Predator-induced phenotypic plasticity of shape and behavior: Parallel and unique patterns across sexes and species. Current Zoology, 63(4). 10.1093/CZ/ZOW072

Atkinson, D. (1994). Temperature and organism size—a biological law for ectotherms? Advances in Ecological Research, 25(C), 1–58. 10.1016/S0065-2504(08)60212-3

Atkinson, D. (1995). Effects of temperature on the size of aquatic ectotherms: Exceptions to the general rule. Journal of Thermal Biology, 20(1–2), 61–74. 10.1016/0306-4565(94)00028-H

Beck, C. W., & Congdon, J. D. (2000). Effects of age and size at metamorphosis on performance and metabolic rates of Southern toad, Bufo terrestris, metamorphs. Functional Ecology, 14(1), 32–38. 10.1046/J.1365-2435.2000.00386.X

Bennett, A. M., Pereira, D., & Murray, D. L. (2013). Investment into defensive traits by anuran prey (Lithobates pipiens) is mediated by the starvation-predation risk trade-off. PLOS ONE, 8(12), e82344. 10.1371/JOURNAL.PONE.0082344

Blanckenhorn, W. U. (1998). Adaptive phenotypic plasticity in growth, development, and body size in the yellow dung fly. Evolution, 52(5), 1394–1407. 10.1111/j.1558-5646.1998.tb02021.x

Boone, M. D. (2005). Juvenile frogs compensate for small metamorph size with terrestrial growth: overcoming the effects of larval density and insecticide exposure. Journal of Herpetology, 39(3), 416–423. 10.1670/187-04A.1

Bourdeau, P. E., & Johansson, F. (2012). Predator-induced morphological defences as by-products of prey behavior: A review and prospectus. Oikos, 121(8), 1175–1190. 10.1111/j.1600-0706.2012.20235.x

Brown, J. H., Gillooly, J. F., Allen, A. P., Savage, V. M., & West, G. B. (2004). Toward a Metabolic Theory of Ecology. Ecology, 85(7), 1771–1789. 10.1890/03-9000

Burkhead, S. E. M., Streid, C. S., Wright, J. T., Bristow, S. A., Hoang, P. L., Oettle, J. L., Pham, A., Pulliam, S., Mardis, D. R., Stybr, E. A., Ward, K. J., & Luhring, T. M. (2022). Aquatic systems of the Wichita State University Biological Field Station: Youngmeyer Ranch, Elk County, Kansas. Transactions of the Kansas Academy of Science, 125(3–4). 10.1660/062.125.0301

Burraco, P., Duarte, L. J., & Gomez-Mestre, I. (2013). Predator-induced physiological responses in tadpoles challenged with herbicide pollution. Current Zoology, 59(4), 475–484. 10.1093/CZOOLO/59.4.475

Buskirk, J. V. (2000). The costs of an inducible defense in anuran larvae. Ecology, 81(10), 2813. 10.2307/177343

Cabrera-Guzmán, E., Crossland, M. R., Brown, G. P., & Shine, R. (2013). Larger body size at metamorphosis enhances survival, growth and performance of young Cane toads (Rhinella marina). PLoS ONE, 8(7), 70121. 10.1371/JOURNAL.PONE.0070121

Calder, W. A. (1984). Size, function, and life history. Harvard University Press.

Calhoun, A. J. K., & DeMaynadier, P. G. (2007). Science and conservation of vernal pools in northeastern North America: ecology and conservation of seasonal wetlands in northeastern North America. CRC Press. 10.1201/9781420005394

Child, T., Phillips, B. L., Brown, G. P., & Shine, R. (2008). The spatial ecology of cane toads (Bufo marinus) in tropical Australia: Why do metamorph toads stay near the water? Austral Ecology, 33(5), 630–640. 10.1111/j.1442-9993.2007.01829.x

Costa, Z. J., & Kishida, O. (2015). Nonadditive impacts of temperature and basal resource availability on predator–prey interactions and phenotypes. Oecologia, 178(4), 1215–1225. 10.1007/S00442-015-3302-X

Courtney Jones, S. K., Munn, A. J., Penman, T. D., & Byrne, P. G. (2015). Long-term changes in food availability mediate the effects of temperature on growth, development and survival in striped marsh frog larvae: Implications for captive breeding programmes. Conservation Physiology, 3(1). 10.1093/CONPHYS/COV029

Crawford, B. A., Hickman, C. R., & Luhring, T. M. (2012). Testing the Threat-Sensitive Hypothesis with predator familiarity and dietary specificity. Ethology, 118(1), 41–48. 10.1111/j.1439-0310.2011.01983.x

Crowl, T. A., & Covich, A. P. (1990). Predator-induced life-history shifts in a freshwater snail. Science, 247(4945), 949–951. 10.1126/science.247.4945.949

Dastansara, N., Vaissi, S., Mosavi, J., & Sharifi, M. (2017). Impacts of temperature on growth, development and survival of larval Bufo (Pseudepidalea) viridis (Amphibia: Anura): Implications of climate change. Zoology and Ecology, 27(3–4), 228–234. 10.1080/21658005.2017.1360037

DeLong, J. P., Gibert, J. P., Luhring, T. M., Bachman, G., Reed, B., Neyer, A., & Montooth, K. L. (2017). The combined effects of reactant kinetics and enzyme stability explain the temperature dependence of metabolic rates. Ecology and Evolution, 7(11), 3940–3950. 10.1002/ece3.2955

Earhart, M. L., Blanchard, T. S., Harman, A. A., & Schulte, P. M. (2022). Hypoxia and high temperature as interacting stressors: will plasticity promote resilience of fishes in a changing world? Biological Bulletin, 243(2), 149–170. 10.1086/722115

Forster, J., & Hirst, A. G. (2012). The temperature-size rule emerges from ontogenetic differences between growth and development rates. Functional Ecology, 26(2), 483–492. 10.1111/j.1365-2435.2011.01958.x

Forster, J., Hirst, A. G., & Atkinson, D. (2012). Warming-induced reductions in body size are greater in aquatic than terrestrial species. Proceedings of the National Academy of Sciences, 109(47), 19310–19314. 10.1073/pnas.1210460109

Forster, J., Hirst, A. G., & Woodward, G. (2011). Growth and development rates have different thermal responses. The American Naturalist, 178(5), 668–678. 10.1086/662174

Frede, T. T., & Lignell, R. (1997). Theoretical models for the control of bacterial growth rate, abundance, diversity and carbon demand. Aquatic Microbial Ecology, 13, 19–27.

Gao, J., Zhang, W., Dang, W., Mou, Y., Gao, Y., Sun, B.-J., & Du, W.-G. (2014). Heat shock protein expression enhances heat tolerance of reptile embryos. Proceedings of the Royal Society B: Biological Sciences, 281(1791), 20141135. 10.1098/rspb.2014.1135

Gherardi, F., Renai, B., & Corti, C. (2001). Crayfish predation on tadpoles: A comparison between a native (Austropotamobius pallipes) and an alien species (Procambarus clarkii). BFPP - Bulletin Francais de La Peche et de La Protection Des Milieux Aquatiques, (361), 659–668. 10.1051/KMAE:2001011

Gibbons, J. W., Winne, C. T., Scott, D. E., Willson, J. D., Glaudas, X., Andrews, K. M., Todd, B. D., Fedewa, L. A., Wilkinson, L., Tsaliagos, R. N., Harper, S. J., Greene, J. L., Tuberville, T. D., Metts, B. S., Dorcas, M. E., Nestor, J. P., Young, C. A., Akre, T., Reed, R. N., … Rothermel, B. B. (2006). Remarkable amphibian biomass and abundance in an isolated wetland: implications for wetland conservation. Conservation Biology, 20(5), 1457–1465. 10.1111/j.1523-1739.2006.00443.x

Gibert, J. P. (2016). The joint effect of phenotypic variation and temperature on predator-prey interactions [Doctoral Dissertation] University of Nebraska.

Gillooly, J. F., Charnov, E. L., West, G. B., Savage, V. M., & Brown, J. H. (2002). Effects of size and temperature on developmental time. Nature, 417(6884), 70–73. 10.1038/417070a

Giovanelli, J. G. R., Haddad, C. F. B., & Alexandrino, J. (2008). Predicting the potential distribution of the alien invasive American bullfrog (Lithobates catesbeianus) in Brazil. Biological Invasions, 10(5), 585–590. 10.1007/BF01875448

Glazier, D. S., & Calow, P. (1992). Energy allocation rules in Daphnia magna: Clonal and age differences in the effects of food limitation. Oecologia, 90(4), 540–549.

Gosner, K. L. (1960). Herpetologists’ League a simplified table for staging anuran embryos and larvae with notes on identification a simplified table for staging anuran embryos and larvae with notes on identification. Herpetologica, 16(23), 183–190. 10.2307/3890061

Grigaltchik, V. S., Webb, C., & Seebacher, F. (2016). Temperature modulates the effects of predation and competition on mosquito larvae. Ecological Entomology, 41(6), 668–675. 10.1111/een.12339

Grosjean, S. (2005). The choice of external morphological characters and developmental stages for tadpole-based anuran taxonomy: A case study in Rana (Sylvirana) nigrovittata (Blyth, 1855) (Amphibia, Anura, Ranidae). Contributions to Zoology, 74(1–2), 61–76. 10.1163/18759866-0740102005

Harkey, G. A., & Semlitsch, R. D. (1988). Effects of temperature on growth, development, and color polymorphism in the ornate chorus frog Pseudacris ornata. Copeia, 1988(4), 1001. 10.2307/1445724

Houseman, G. R., Kraushar, M. S., & Rogers, C. M. (2016). The Wichita State University biological field station: bringing breadth to research along the precipitation gradient in kansas. Kansas Academy of Science, 119(1), 27–32. 10.1660/062.119.0106

Howe, N. R. (1976). Behavior of sea anemones evoked by the alarm pheromone anthopleurine. Journal of Comparative Physiology, 107(1), 67–76. 10.1007/BF00663919

Huey, R. B., & Kingsolver, J. G. (2019). Climate warming, resource availability, and the metabolic meltdown of ectotherms. American Naturalist, 194(6), E140–E150. 10.1086/705679

Katzenberger, M., Hammond, J., Duarte, H., Tejedo, M., Calabuig, C., & Relyea, R. A. (2014). Swimming with predators and pesticides: How environmental stressors affect the thermal physiology of tadpoles. PLoS ONE, 9(5). 10.1371/journal.pone.0098265

Lent, E. M., & Babbitt, K. J. (2020). The effects of hydroperiod and predator density on growth, development, and morphology of wood frogs (Rana sylvatica). Aquatic Ecology, 54(1), 369–386. 10.1007/s10452-020-09748-y

Lowe, W. H., Martin, T. E., Skelly, D. K., & Woods, H. A. (2021). Metamorphosis in an era of increasing climate variability. Trends in Ecology & Evolution, 36(4), 360–375. 10.1016/j.tree.2020.11.012

Luhring, T. M., & Delong, J. P. (2016). Predation changes the shape of thermal performance curves for population growth rate. Current Zoology, 62(5), 501–505. 10.1093/cz/zow045

Luhring, T. M., Vavra, J. M., Cressler, C. E., & DeLong, J. P. (2018). Predators modify the temperature dependence of life-history trade-offs. Ecology and Evolution, 8(17), 8818–8830. 10.1002/ece3.4381

Luhring, T. M., Vavra, J. M., Cressler, C. E., & DeLong, J. P. (2019). Phenotypically plastic responses to predation risk are temperature dependent. Oecologia, 191(3), 709–719. 10.1007/s00442-019-04523-9

Maher, J. M., Werner, E. E., & Denver, R. J. (2013). Stress hormones mediate predator-induced phenotypic plasticity in amphibian tadpoles. Proceedings of the Royal Society B: Biological Sciences, 280(1758). 10.1098/RSPB.2012.3075

Masson-Delmotte, V., Zhai, P., Pirani, A., Connors, S. L., Péan, C., Berger, S., Caud, N., Chen, Y., Goldfarb, L., & Gomis, M. I. (2021). Climate change 2021: The physical science basis. Contribution of Working Group I to the Sixth Assessment Report of the Intergovernmental Panel on Climate Change, 2(1), 2391.

Mauro, A. A., Shah, A. A., Martin, P. R., & Ghalambor, C. K. (2022). An integrative perspective on the mechanistic basis of context- dependent species interactions. Integrative and Comparative Biology, 62(2), 164–178. 10.1093/ICB/ICAC055

McPeek, M. A., Grace, M., & Richardson, J. M. L. (2001). Physiological and behavioral responses to predators shape the growth/predation risk trade-off in damselflies. Ecology, 82(6), 1535–1545. https://doi-org.proxy2.cl.msu.edu/10.1890/0012-9658(2001)082[1535:PABRTP]2.0.CO;2

Mishra, P., Naik, S., Babu, P. V., Pradhan, U., Begum, M., Kaviarasan, T., Vashi, A., Bandyopadhyay, D., Ezhilarasan, P., Panda, U. S., & Murthy, M. V. R. (2022). Algal bloom, hypoxia, and mass fish kill events in the backwaters of Puducherry, Southeast coast of India. Oceanologia, 64(2), 396–403. 10.1016/J.OCEANO.2021.11.003

Monaghan, P. (2008). Early growth conditions, phenotypic development and environmental change. Philosophical Transactions of the Royal Society B: Biological Sciences, 363(1497), 1635. 10.1098/RSTB.2007.0011

Mori, T., Hiraka, I., Kurata, Y., Kawachi, H., Kishida, O., & Nishimura, K. (2005). Genetic basis of phenotypic plasticity for predator-induced morphological defenses in anuran tadpole, Rana pirica, using cDNA subtraction and microarray analysis. Biochemical and Biophysical Research Communications, 330(4), 1138–1145. 10.1016/J.BBRC.2005.03.091

National Research Council. (1974). Amphibians: Guidelines for the Breeding, Care and Management of Laboratory Animals. The National Academies Press. Washington, DC. 10.17226/661

Newman, R. A. (1998). Ecological constraints on amphibian metamorphosis: Interactions of temperature and larval density with responses to changing food level. Oecologia 1998 115:1, 115(1), 9–16. 10.1007/S004420050485

Ohmer, M. E. B., Hammond, T. T., Switzer, S., Wantman, T., Bednark, J. G., Paciotta, E., Coscia, J., & Richards-Zawacki, C. L. (2023). Developmental environment has lasting effects on amphibian post-metamorphic behavior and thermal physiology. Journal of Experimental Biology, 226(9). 10.1242/jeb.244883

Padfield, D., Castledine, M., & Buckling, A. (2020). Temperature-dependent changes to host–parasite interactions alter the thermal performance of a bacterial host. ISME Journal, 14(2), 389–398. 10.1038/s41396-019-0526-5

Pascual, L. S., Segarra-Medina, C., Gómez-Cadenas, A., López-Climent, M. F., Vives-Peris, V., & Zandalinas, S. I. (2022). Climate change-associated multifactorial stress combination: A present challenge for our ecosystems. Journal of Plant Physiology, 276, 153764. 10.1016/j.jplph.2022.153764

Pauwels, K., Stoks, R., & De Meester, L. (2005). Coping with predator stress: Interclonal differences in induction of heat-shock proteins in the water flea Daphnia magna. Journal of Evolutionary Biology, 18(4), 867–872. 10.1111/J.1420-9101.2005.00890.X

Perrin, N. (1995). About Berrigan and Charnov Life-History Puzzle. OIKOS, 73(1), 137–139. 10.2307/3545737

Peters, R. H. (1983). The Ecological Implications of Body Size. Cambridge University Press. 10.1017/CBO9780511608551

Polo-Cavia, N., Boyero, L., Martín-Beyer, B., Barmuta, L. A., & Bosch, J. (2017). Joint effects of rising temperature and the presence of introduced predatory fish on montane amphibian populations. Animal Conservation, 20(2), 128–134. 10.1111/acv.12294

Rebolledo, A. P., Sgrò, C. M., & Monro, K. (2021). Thermal Performance Curves Are Shaped by Prior Thermal Environment in Early Life. Frontiers in Physiology, 12, 738338. 10.3389/FPHYS.2021.738338

Relyea, R. A. (2002). Competitor-Induced Plasticity in Tadpoles: Consequences, Cues, and Connections to Predator-Induced Plasticity. Ecological Monographs, 72(4), 523–540. https://doi-org.proxy2.cl.msu.edu/10.1890/0012-9615(2002)072[0523:CIPITC]2.0.CO;2

Relyea, R. A. (2006). The effects of pesticides, pH, and predatory stress on amphibians under mesocosm conditions. Ecotoxicology, 15(6), 503–511. 10.1007/S10646-006-0086-0

Relyea, R. A. (2007). Getting out alive: How predators affect the decision to metamorphose. Oecologia, 152(3), 389–400. 10.1007/s00442-007-0675-5

Relyea, R. A., & Hoverman, J. T. (2003). The impact of larval predators and competitors on the morphology and fitness of juvenile treefrogs. Oecologia, 134(4), 596–604. 10.1007/s00442-002-1161-8

Rezende, E. L., & Bozinovic, F. (2019). Thermal performance across levels of biological organization. Philosophical Transactions of the Royal Society B, 374(1778). 10.1098/RSTB.2018.0549

Riessen, H. P. (1999). Predator-induced life history shifts in Daphnia: A synthesis of studies using meta-analysis. Canadian Journal of Fisheries and Aquatic Sciences, 56(12), 2487–2494. 10.1139/f99-155

Riessen, H. P., & Gilbert, J. J. (2019). Divergent developmental patterns of induced morphological defenses in rotifers and Daphnia: Ecological and evolutionary context. Limnology and Oceanography, 64(2), 541–557. 10.1002/lno.11058

Rita, A., Cerbo, D., & Biancardi, C. M. (2010). Morphometric study on tadpoles of Bombina variegata. Acta Herpetologica, 5(2), 223–232. 10.13128/ACTA_HERPETOL-9036

Saeed, M., Rais, M., Gray, R. J., Ahmed, W., Akram, A., Gill, S., & Fareed, G. (2021). Rise in temperature causes decreased fitness and higher extinction risks in endemic frogs at high altitude forested wetlands in northern Pakistan. Journal of Thermal Biology, 95, 102809. 10.1016/J.JTHERBIO.2020.102809

Schneider, C. A., Rasband, W. S., & Eliceiri, K. W. (2012). NIH Image to ImageJ: 25 years of image analysis. Nature Methods, 9(7), 671–675. 10.1038/nmeth.2089

Schoeppner, N. M., & Relyea, R. A. (2005). Damage, digestion, and defence: The roles of alarm cues and kairomones for inducing prey defences. Ecology Letters, 8(5), 505–512. 10.1111/J.1461-0248.2005.00744.X

Schoeppner, N. M., & Relyea, R. A. (2009). Interpreting the smells of predation: How alarm cues and kairomones induce different prey defences. Functional Ecology, 23(6), 1114–1121. 10.1111/j.1365-2435.2009.01578.x

Schulte, P. M. (2015). The effects of temperature on aerobic metabolism: Towards a mechanistic understanding of the responses of ectotherms to a changing environment. Journal of Experimental Biology, 218(12), 1856–1866. 10.1242/jeb.118851

Semlitsch, R. D., & Reyer, H.-U. (1992). Modification of anti-predator behaviour in tadpoles by environmental conditioning. Journal of Animal Ecology, 61, 353–360.

Sinsch, U., Leus, F., Sonntag, M., & Hantzschmann, A. M. (2020). Carry-over effects of the larval environment on the post-metamorphic performance of Bombina variegata (Amphibia, Anura). Herpetological Journal, 30(3), 126–134. 10.33256/HJ30.3.126134

Skelly, D. K., & Werner, E. E. (1990). Behavioral and Life-Historical Responses of Larval American Toads to an Odonate Predator. Ecology, 71(6), 2313–2322. 10.2307/1938642

Skerlec, S., & Luhring, T. M. (2023). Investigating the impacts of drought-related drying events following pond refill on the growth, survival, and postmetamorphic fitness of Lithobates blairi tadpoles [Master’s thesis]. Wichita State University.

Slos, S., & Stoks, R. (2008). Predation risk induces stress proteins and reduces antioxidant defense. Functional Ecology, 22(4), 637–642. 10.1111/J.1365-2435.2008.01424.X

Sokolova, I. M., & Francisco, S. (2013). Energy-Limited Tolerance to Stress as a Conceptual Framework to Integrate the Effects of Multiple Stressors. Integrative and Comparative Biology, 53(4), 597–608. 10.1093/ICB/ICT028

Sørensen, J. G., Pekkonen, M., Lindgren, B., Loeschcke, V., Laurila, A., & Merilä, J. (2009). Complex patterns of geographic variation in heat tolerance and Hsp70 expression levels in the common frog Rana temporaria. Journal of Thermal Biology, 34(1), 49–54. 10.1016/J.JTHERBIO.2008.10.004

Steiner, U. K. (2007). Investment in defense and cost of predator-induced defense along a resource gradient. Oecologia, 152(2), 201–210. 10.1007/S00442-006-0645-3

Steiner, U. K., & Van Buskirk, J. (2008). Environmental stress and the costs of whole-organism phenotypic plasticity in tadpoles. Journal of Evolutionary Biology, 21(1), 97–103. 10.1111/J.1420-9101.2007.01463.X

Steinwascher, K. (1981). Competition for two resources. Oecologia, 49(3), 415–418. 10.1007/BF00347609

Székely, D., Cogălniceanu, D., Székely, P., Armijos-Ojeda, D., Espinosa-Mogrovejo, V., & Denoël, M. (2020). How to recover from a bad start: Size at metamorphosis affects growth and survival in a tropical amphibian. BMC Ecology, 20(1), 24. 10.1186/s12898-020-00291-w

Thomas, M. K., Aranguren-Gassis, M., Kremer, C. T., Gould, M. R., Anderson, K., Klausmeier, C. A., & Litchman, E. (2017). Temperature–nutrient interactions exacerbate sensitivity to warming in phytoplankton. Global Change Biology, 23(8), 3269–3280. 10.1111/gcb.13641

Thompson, C. M., & Popescu, V. D. (2021). Complex hydroperiod induced carryover responses for survival, growth, and endurance of a pond-breeding amphibian. Oecologia, 195(4), 1071–1081. 10.1007/s00442-021-04881-3

Travis, J. (1984). Anuran Size at Metamorphosis: Experimental Test of a Model Based on Intraspecific Competition. Ecology, 65(4), 1155–1160. 10.2307/1938323

Trussell, G. C., Ewanchuk, P. J., & Matassa, C. M. (2006). The Fear of Being Eaten Reduces Energy Transfer in a Simple Food Chain. Ecology, 87(12), 2979–2984. https://doi-org.proxy2.cl.msu.edu/10.1890/0012-9658(2006)87[2979:TFOBER]2.0.CO;2

Urban, M. C. (2008). Salamander evolution across a latitudinal cline in gape-limited predation risk. Oikos, 117(7), 1037–1049. 10.1111/J.0030-1299.2008.16334.X

Urban, M. C., Richardson, J. L., Freidenfelds, N. A., Drake, D. L., Fischer, J. F., & Saunders, P. P. (2017). Microgeographic Adaptation of Wood Frog Tadpoles to an Apex Predator. Copeia, 105(3), 451–461. 10.1643/CG-16-534

Ward, K. J., Burkhead, S. E. M., Stybr-Burrus, E. A., & Luhring, T. M. (2023). No free refills: Prior drying and vertebrate colonisation alter ecological functioning and vertebrate fitness within experimental aquatic systems. Freshwater Biology. 10.1111/fwb.14157

Werner, E. E. (1986). Amphibian Metamorphosis: Growth Rate, Predation Risk, and the Optimal Size at Transformation. The American Naturalist, 128(3), 319–341. 10.1086/284565

White, E. P., Ernest, S. K. M., Kerkhoff, A. J., & Enquist, B. J. (2007). Relationships between body size and abundance in ecology. Trends in Ecology & Evolution, 22(6), 323–330. 10.1016/j.tree.2007.03.007

Wilbur, H. M. (1980). Complex Life Cycles. Annual Review of Ecology and Systematics, 11, 67–93. 10.1146/annurev.es.11.110180.000435

Williams, I. N., Torn, M. S., Riley, W. J., & Wehner, M. F. (2014). Impacts of climate extremes on gross primary production under global warming. Environmental Research Letters, 9(9), 094011. 10.1088/1748-9326/9/9/094011

Wood, S.N. (2006) Generalized Additive Models: An Introduction with R. CRC Press, Boca Raton, FL.

Zedler, P. H. (2003). Vernal pools and the concept of “isolated wetlands.” Wetlands, 23, 597–607.

Zhu, W., Zhao, T., Zhao, C., Li, C., Xie, F., Liu, J., & Jiang, J. (2023). How will warming affect the growth and body size of the largest extant amphibian? More than the temperature–size rule. Science of the Total Environment, 859. 10.1016/J.SCITOTENV.2022.160105

